# A method for RNA structure prediction shows evidence for structure in lncRNAs

**DOI:** 10.1101/284869

**Authors:** Riccardo Delli ponti, Alexandros Armaos, Stefanie Marti, Gian Gaetano Tartaglia

## Abstract

To compare the secondary structures of RNA molecules we developed the *CROSSalign* method. *CROSSalign* is based on the combination of the Computational Recognition Of Secondary Structure (CROSS) algorithm to predict the RNA secondary structure at single-nucleotide resolution using sequence information, and the Dynamic Time Warping (DTW) method to align profiles of different lengths. We applied *CROSSalign* to investigate the structural conservation of long non-coding RNAs such as XIST and HOTAIR as well as ssRNA viruses including HIV. In a pool of sequences with the same secondary structure *CROSSalign* accurately recognizes repeat A of XIST and domain D2 of HOTAIR and outperforms other methods based on covariance modelling. *CROSSalign* can be applied to perform pair-wise comparisons and is able to find homologues between thousands of matches identifying the exact regions of similarity between profiles of different lengths. The algorithm is freely available at the webpage http://service.tartaglialab.com//new_submission/CROSSalign.

## Introduction

Sequence similarity is often considered the key feature to investigate evolutionary conservation of coding transcripts [1]. Yet, knowledge of secondary structure provides important insights into the biological function of RNAs by allowing the study of physical properties, such as for instance molecular interactions [2]. In most cases, information about the RNA folding complements sequence analysis [3] and is useful to understand mechanisms of action: microRNA precursors, for example, are processed by DGCR8 only if properly folded in hairpin loop structures [4]. Similarly, the architecture of ribosomal RNAs evolves in a self-contained way through conservation of stem loops present in ancient species [5,6], indicating distinct requirements for structural elements.

Long non-coding RNAs (lncRNAs) are regarded as a mystery in terms of sequence and structural conservation [7]. The vast majority of lncRNAs evolve under little or no selective constraints, undergo almost no purifying selection, show poor expression, and do not have often easily identifiable orthologues [7,8]. Indeed, the average sequence homology of evolutionarily conserved lncRNAs is 20% between human and mouse and drops to 5% between human and fish [7]. Thus, primary structure does not provide relevant information to study lncRNA conservation and secondary structure could be used for better characterization. Similarly to lncRNAs, the transcriptomes of single-stranded RNA (ssRNA) viruses retain their fold even if sequences mutate rapidly [9], which indicates that secondary structure investigation could be key to revealing evolutionary properties.

To study the structural conservation of two RNA molecules, we developed the *CROSSalign* method. *CROSSalign*, available at our webpages http://service.tartaglialab.com//new_submission/CROSSalign, is based on the combination of two methods: 1) Computational Recognition Of Secondary Structure (CROSS), which is an algorithm trained on experimental data to predict RNA secondary structure profiles without sequence length restrictions and at singlenucleotide resolution [10]; 2) the Dynamic Time Warping (DTW) algorithm to assess the similarity of two profiles of different lengths [11]. DTW flexibility allows managing profiles of different lengths without having to sacrifice computational time.

We applied *CROSSalign* on lncRNAs of different species, compared it with covariation models, as well as on ssRNA viruses. *CROSSalign* is able to find structural homologues among millions of possible matches identifying structural domains with great accuracy.

## Results

To test the performances and functionality of CROSS combined with DTW (**Supplementary Figure. 1** and **2**), we selected a dataset of 22 structures for which crystallographic (exact base pairing between nucleotides) and Selective 2’-hydroxyl acylation analyzed by primer extension (SHAPE; chemical probing of flexible regions used to assess whether a nucleotide is in double- or single-stranded state) data are available [12]. Using DTW, we calculated the structural distances between all possible pairs in the dataset considering crystallographic (dots and parentheses were transformed into binary code) data as well as 1) SHAPE profiles (Area Under the ROC Curve AUC of 0.76, Positive Predictive Value PPV of 0.76 when compared to crystallographic data) and 2) CROSS profiles (AUC 0.72, PPV 0.74 when compared to crystallographic data, see also http://service.tartaglialab.com/staticfiles/algorithms/cross/documentation.html#5).

Structural distances between CROSS profiles show higher correlations to the distances between the respective profiles from crystallographic data (Pearson’s correlation of 0.91) than structural distances between SHAPE profiles (**Figure 1A, 1B**, correlation of 0.50). Moreover, CROSS shows better performances than algorithms such as RNAstructure [13,14] and RNAfold [12] (respective correlations 0.71 and 0.47 with crystals; **Supplementary Figure. 3** and **4**).

Sequence similarity analysis (computed with EMBOSS; see **Material and Methods**) shows comparable correlations with structural distances calculated with either CROSS or crystallographic profiles (respectively: 0.80, 0.83, 0.38 with crystallographic, CROSS and SHAPE data). While CROSS and crystallographic profiles identify specific clusters of RNA molecules with low sequence identity and high structural similarity (colored in red, orange and green according to difference confidence thresholds; **Figure. 1C, 1D, 1E**), SHAPE data cannot be used to identify these structures.

**Figure 1.**
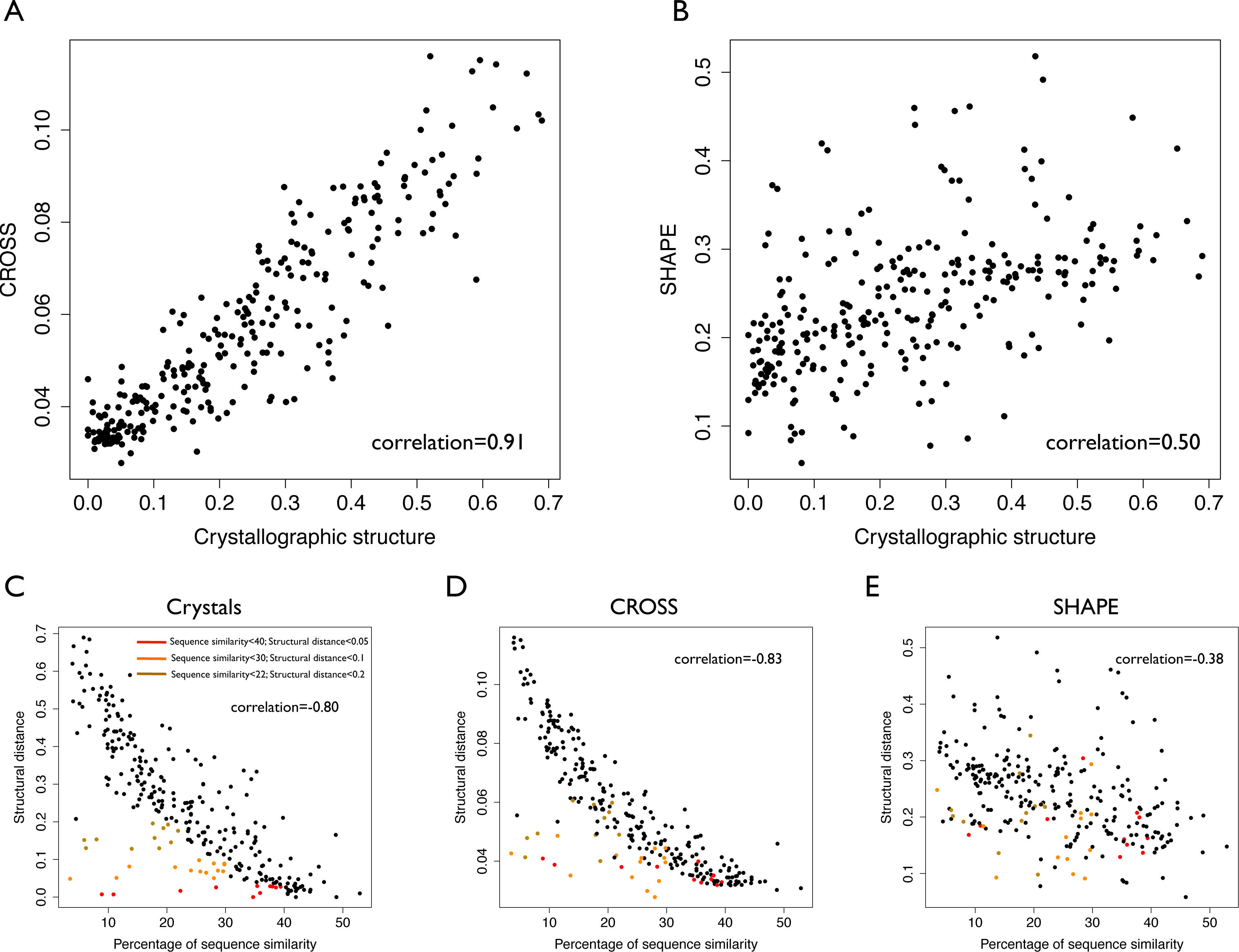
Validation of the CROSSalign method. (**A**) Structural distances correlation between CROSS and crystallographic profiles computed on 22 structures (standard-DTW). (**B**) Correlation between structural distances computed with SHAPE and crystallographic data on the same data (standard-DTW). (**C**) Correlation between structural distances (crystallographic profiles) and sequence similarity. Clusters of similar structures and different sequences (sequence similarity < 40%) are highlighted in brown (structural score < 0.2), orange (structural score < 0.1) and (red structural score < 0.05). (**D**) Correlation between structural distances (CROSS) and sequence similarity. The clusters previously identified for crystallographic data are shown in the plot. (**E**) Correlation between structural distances (**SHAPE**) and sequence similarity. In this case the clusters previously identified are disrupted.

To further test the usefulness of *CROSSalign*, we compared its output with that of CMfmder [15]. CMfmder is a method to compute multiple sequence alignments that exploits structural motifs for ranking (**Materials and Methods, *Comparisons with CMfinder***). We analyzed the largest multiple sequence alignments reported in the CMfmder test set (cobalamin, intron gp II, s box, lysine and histone 3) and used the minimal structural distance to assign the closest match to each transcript. Selecting equal-size groups (lowest and highest CMfinder scores), we measured *CROSSalign* performances on CMfinder rankings, achieving an AUC of 0.80 (**Supplementary Methods; Supplementary Figure 5A**). We note that *CROSSalign* has particularly strong performances on the largest dataset: cobalamin (71 sequences of 216nt; AUC of 0.95; **Supplementary Figure 5B**).

### Ribosomal RNAs

Ribosomal RNAs are considered one of the most ancient, structured and conserved classes of RNA molecules [5]. The first step to validate *CROSSalign* was to search for structural similarities between the *Small SubUnit* (SSU) of the rRNA of different bacteria (**Pseudomonas aeruginosa, Escherichia coli, Bacillus subtilis, Deinococcus radiodurans**). All the rRNAs show significant structural similarity (p-value <10^−5^) with a low structural distance (~0.08; **Table 1A**). By contrast, shuffling one of the two sequences in *CROSSalign* predictions results in non-significant scoring (p-values of 0.10 or higher; **Table 1A**), indicating the importance of the sequence context in our calculations.

**Table 1.** Conservation of ribosomal structures. (**A**) Above the diagonal: structural distances and p-values of bacterial SSUs; below the diagonal: structural distances and p-values upon shuffling one of the two bacterial sequences (Genbank/NCBI: J01859.1, NR_026078, NR_074411.1, bacillus and NR_102783.2). (**B**) Structural distance and p-values of SSUs against E. coli (Genbank/NCBI: M10098.1, NR_074375.1 and NR_132222.1). The shuffled sequence of Escherichia coli is used as a negative control. A coding human mRNA randomly selected from ENSEMBL with the same length of E. coli ribosome (ENSG00000002933) is used as an additional control to highlight the exclusive structural similarities between the different SSUs.

|  | **B. subtilis** | **D. radiodurans** | **E.coli** | **P. aeruginosa** | **E. coli (shuffled)** |
| --- | --- | --- | --- | --- | --- |
| **B. subtilis** | **0** | **0.086**<br>(p-value<10<sup>-5</sup>) | **0.088**<br>(p-value<10<sup>-5</sup>) | **0.082**<br>(p-value<10<sup>-5</sup>) | 0,098<br>(p-value=0,19) |
| **D. radiodurans** | 0,10<br>(p-value=0,35) | **0** | **0.085**<br>(p-value<10<sup>-5</sup>) | **0.087**<br>(p-value<10<sup>-5</sup>) | 0,10<br>(p-value=0,35) |
| **E. coli** | 0,098<br>(p-value=0,19) | 0,10<br>(p-value=0,35) | **0** | **0.066**<br>(p-value<10<sup>-5</sup>) | 0,098<br>(p-value=0,19) |
| **P. aeruginosa** | 0,097<br>(p-value=0,13) | 0,10<br>(p-value=0,35) | 0,098<br>(p-value=0,19) | **0** | 0,098<br>(p-value=0,19) |

**Table 1. Conservation of ribosomal structures.**
|  | **E. Coli** | **E. coli (shuffled)** | **H. sapiens (mRNA)** |
| --- | --- | --- | --- |
| **H. sapiens** | **0,079**<br>(p-value<10<sup>-5</sup>) | 0,097<br>(p-value=0,13) | 0,096<br>(p-value=0,10) |
| **P. furious** | **0,077**<br>(p-value<10<sup>-5</sup>) | 0,096<br>(p-value=0,10) | 0,096<br>(p-value=0,10) |
| **S. cerevisiae** | **0,079**<br>(p-value<10<sup>-5</sup>) | 0,100<br>(p-value=0,35) | 0,095<br>(p-value=0,05) |

*Pseudomonas aeruginosa and Escherichia coli* are the most similar SSUs (structural distance of 0.06; p-value <10^−5^). As the SSU of the rRNAs is thought to have evolved in a self-contained structure where the secondary structure of the ancient species is contained in the other ones [6], we searched the complete SSU of *Escherichia coli in the SSU of other species such as Pyrococcus furiosus, Saccharomyces cerevisiae and Homo sapiens*. The results show a strong and significant structural similarity, in agreement with the theory of self-contained evolution (**Table 1B**). By contrast, comparison with randomized E. coli SSU or H. sapiens mRNA of the same length results in non-significant scores (p-values of 0.10 or higher **Table 1B**).

### XIST

*XIST* is a lncRNA characterized by several repetitive domains showing different structural properties (**Figure. 2A** and **2B**) [16]. The 5’ conserved region, named A-repeat (or RepA), is indispensable for gene-silencing and has been shown to be highly conserved in mammals. In mouse it consists of 7.5 copies (8.5 in humans) of 26-mers separated by U-rich linkers (**Figure 2A**) [17]. In 2015 and 2017, two reports on the *XIST* A-repeat structure were published [18,19], both making use of experimental techniques to infer *XIST* structure. The A-repeat structures obtained are similar with strikingly identical stem-loop structures, both emerging from larger RNA bulges of repeats 3, 5, and 6 (for a comparison with CROSS predictions, see our previous manuscript [10]).

**Figure 2.**
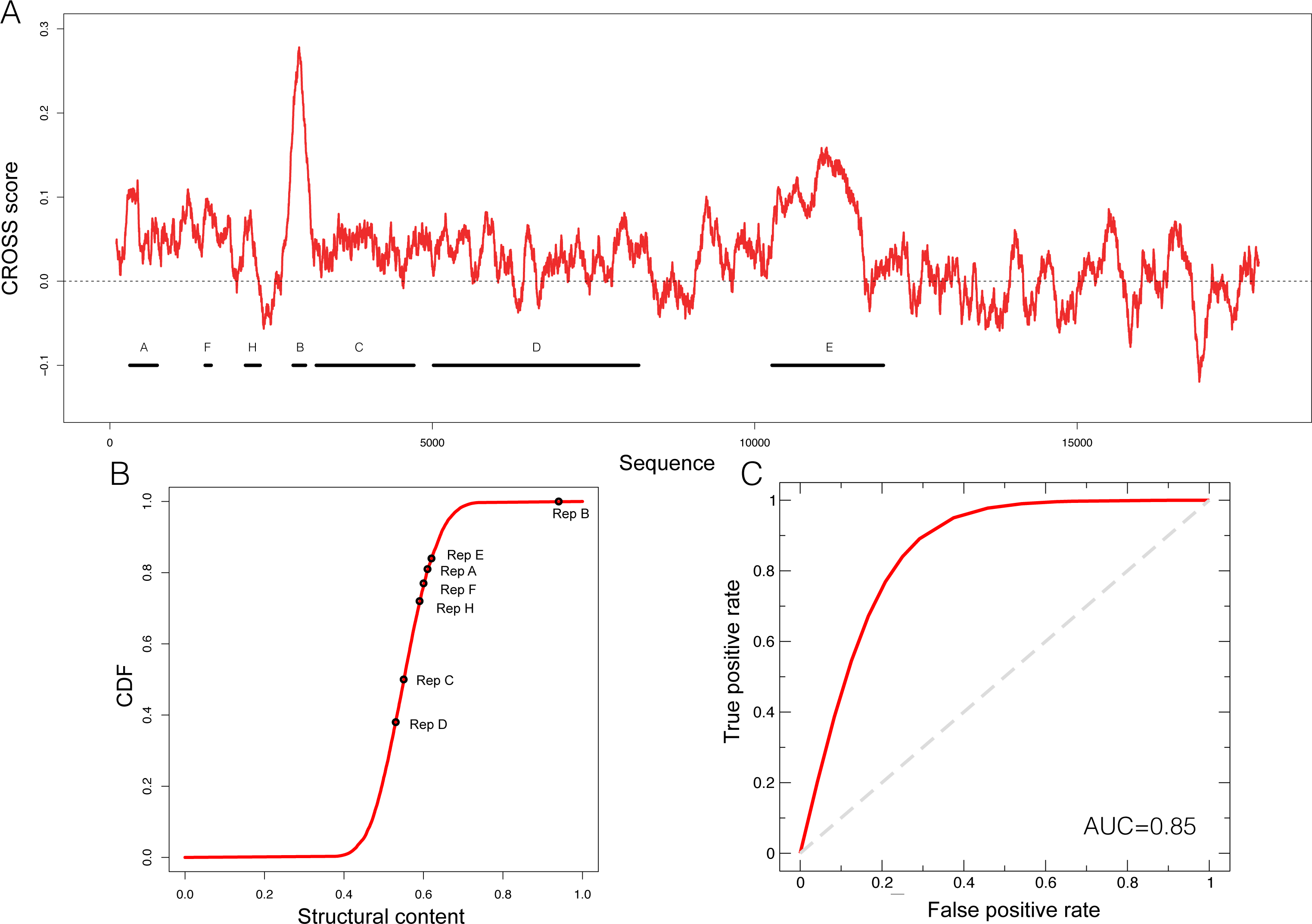
Predictions of XIST structure. (**A**) Secondary structure profile of murine *XIST* obtained using CROSS. Positive regions are to be considered double-stranded, negative regions single-stranded. (**B**) Cumulative distribution function (**CDF**) of the structural content of all the human lincRNAs predicted by CROSS. The structural contents of the Rep domains of *XIST* are reported on the curve. (**C**) ROC curve of *CROSSalign* to identify reverse engineered sequences with the same structure as RepA and a different sequence.

To evaluate the ability of *CROSSalign* to recognize the structural content in RepA, we used *RNAinverse* from Vienna suite [12] to generate 50 different sequences (average identity of 40%; **Supplementary Table 1**) with the same structure as RepA (**Materials and Methods**; ***Reverse engineering: from structure to sequence***). The sequences were then divided into a reference (25 transcripts) and a positive (25 transcripts) set, and we built a list of negative cases by shuffling 25 times the original RepA. We then used *CROSSalign* to compute all scores for the positive and negative set against the corresponding reference set and used the minimal structural distance to assign the closest match to each transcript. The strong performances obtained highlight the ability of *CROSSalign* to identify structural similarities regardless of the sequence similarity (AUC of 0.85; **Figure 2C**).

As a further test of the usefulness of *CROSSalign* we compared the above results with those obtained using *CMsearch* from the Infernal-1.1.2 package, an algorithm based on a covariance model approach [20]. In this case, *CMsearch* is not able to identify any match between either the positive or negative list and the reference sets (AUC of 0.5; **Materials and Methods**; ***Comparisons with CMsearch***). These results indicate that *CROSSalign* is able to recognize structural similarities between non-similar sequences, outperforming covariance-based approaches such as *CMsearch*.

After this first validation step, we used *CROSSalign* to study structural similarities of *XIST* domain RepA in 10 different species [21]. Our analysis reveals that primates cluster close to human (**Supplementary Figure 6A**) while other species are more distant (**Supplementary Figure. 6A** and **7**). By contrast, calculating sequence similarity with respect to human *XIST* (computed with EMBOSS; see **Materials and Methods**), we could not identify a specific cluster for primates (Supplementary Figure 6B). Thus, our results indicate that secondary structure shows a higher degree of conservation than sequence.

We then selected RepA of orangutan and searched for structural similarities within all human intergenic lncRNAs (lincRNAs 8176 sequences; ENSEMBLE 82). *XIST* was ranked as the best significant match in the pool (structural distance 0.01; p-value < 10-6) and RepA was correctly identified (predicted coordinates: 328-764; 95% overlap with the query region; **Figure 3A; Supplementary Table 2**). Similar results were observed for baboon RepA (best result: 0.032; 86% overlap with the query region) and lemur RepA (best result: 0.075; p-value; 97% overlap with the query region), suggesting a strong structural conservation within primates (**Figure 3B**). By contrast, human and mouse RepA showed a larger distance in terms of both structural and sequence similarity, which is in agreement with previous studies on lncRNA conservation [22].

**Figure 3.**
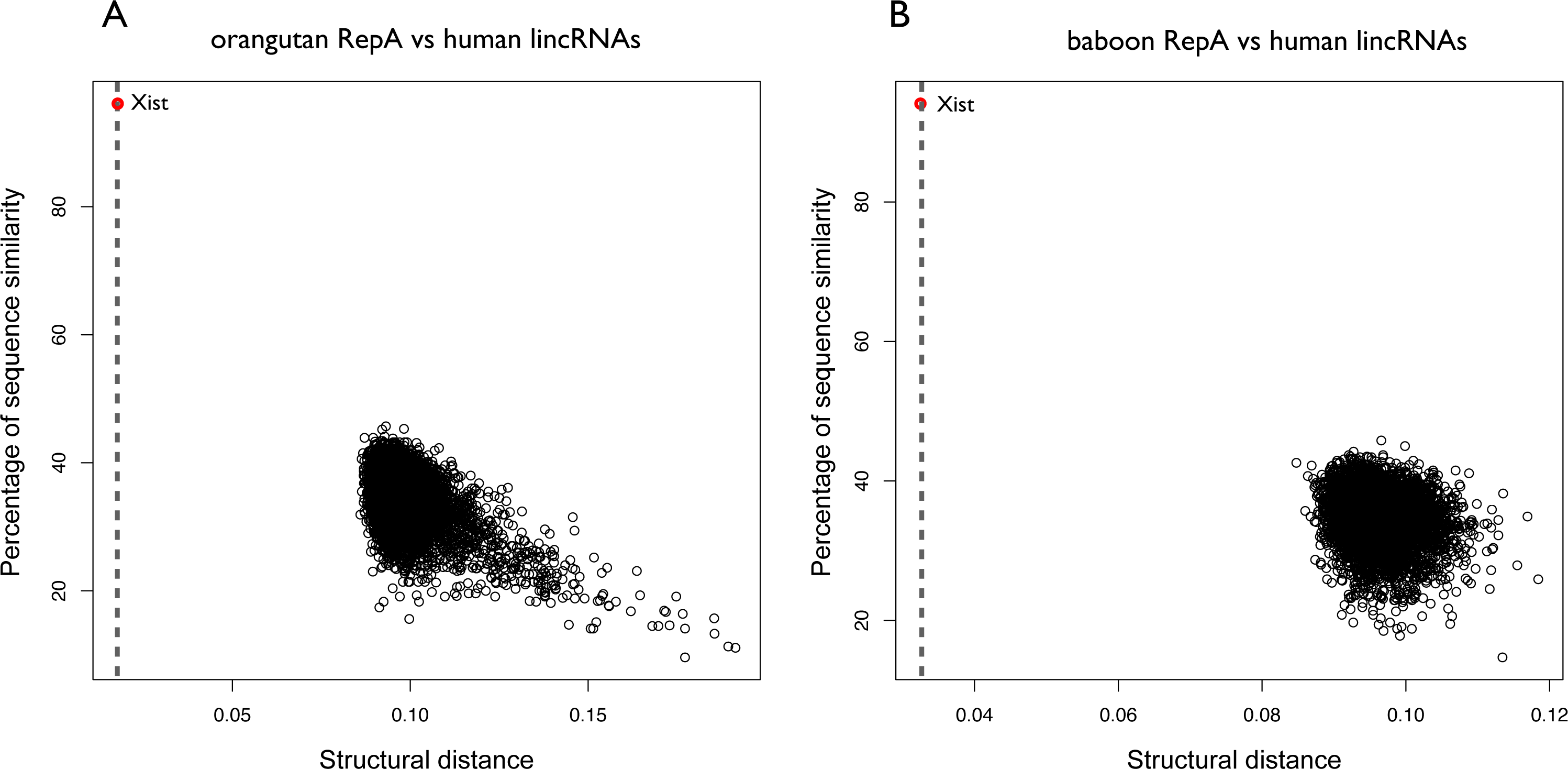
Structural conservation of XIST RepA within primates. (**A**) Structural distances of Orangutan RepA are computed against all human lincRNAs. The structural distance is calculated using OBE-DTW and plotted against sequence similarity. Orangutan RepA is identified as the best match (colored in red). (**B**) Structural distances of baboon RepA against all the lincRNAs of human. Baboon RepA is identified as the best match (colored in red).

We used *CROSSalign* to search for the human RepA within all mouse lncRNAs and identified *XIST* as the 5^th^ best hit (structural distance 0.085; p-value < 10^−6^; Figure 4A). In this case, the position of RepA was not correctly assigned (coordinates: 10306-10698; 0% overlap) but the best match falls into the regulatory region of exon 7, and the structural relation between RepA and exon 7 has been reported [23]. Importantly, the correct coordinates of human RepA within mouse *XIST* rank second in our analysis (structural distance 0.086; p-value < 10^−6^), while the best match is a miRNA-containing the gene Mirg (ENSMUSG00000097391) and the two secondary structure profiles show a strong correlation of 0.92 (**Figure 4B**). Interestingly, even if little information is available on Mirg, the transcript is prevalently expressed in the embryo [24]. This result unveils a previously unreported relationship between *XIST* and Mirg, in which structural and functional homologies can be linked. Intriguingly, also the second best result, Rian (ENSMUSG00000097451), is expressed in embryo, while no information is available on the third and fourth hits (ENSMUSG00000107391 and ENSMUSG00000085312). We note that the five matches here are not listed in the top 20 hits obtained by analysis of sequence similarity (<34%).

**Figure 4.**
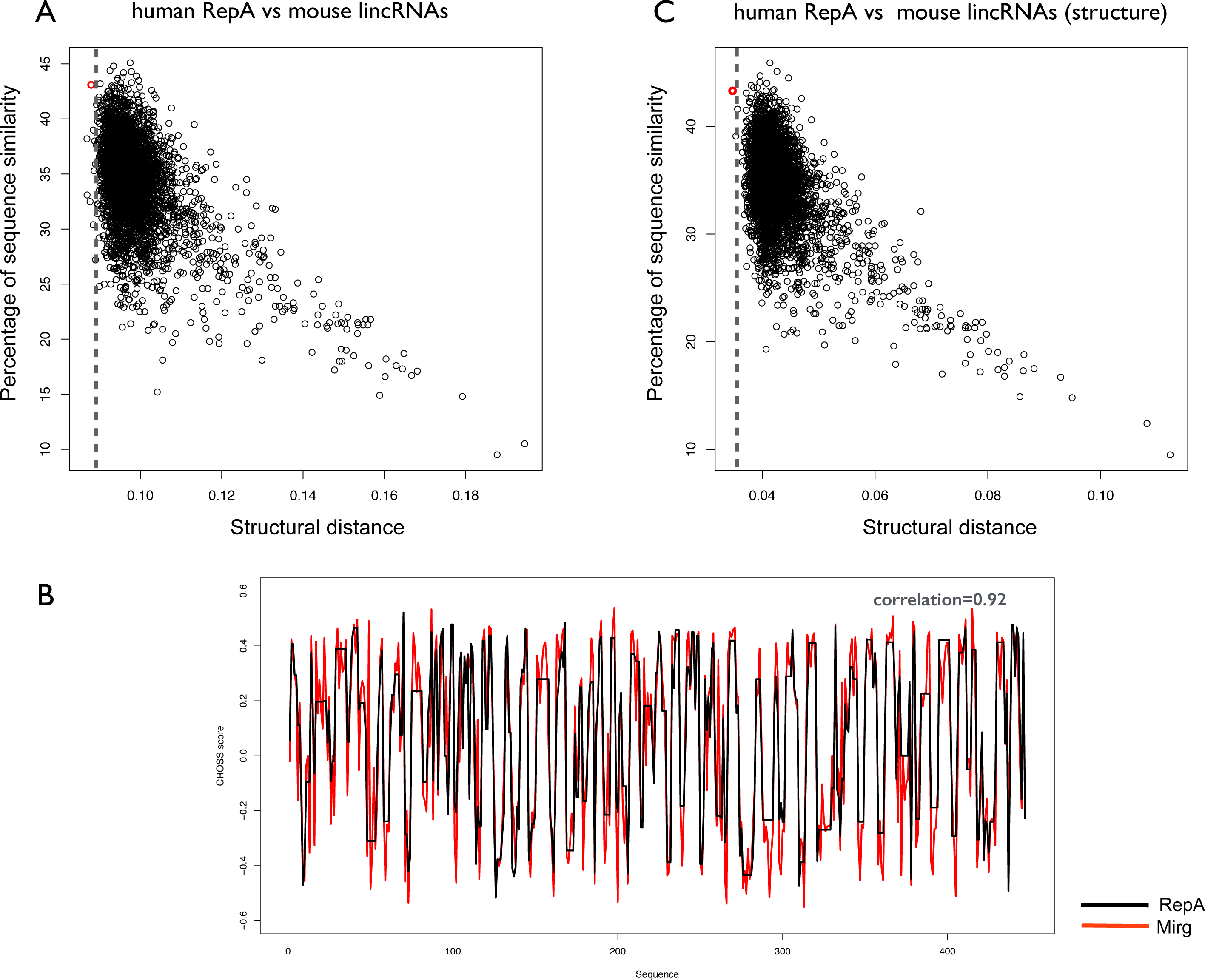
Structural similarities between human and mouse XIST RepA. (**A**) Structural similarities of human RepA against all mouse lincRNAs. The structural distance was calculated using *CROSSalign* (OBE-DTW) and plotted against sequence similarity. Human RepA is not the best match (5th best hit; colored in red). (**B**) Structural similarities of human RepA against all mouse lincRNAs using doublestranded nucleotides (nucleotide with CROSS score < 0 are set to 0). Human RepA is identified as the best match (colored in red), which highlights the importance of the structural content for the regulatory domains of the lncRNAs. (**C**) Secondary structure profile of human RepA, obtained as optimal path with OBE-DTW, compared with the best match in mouse lincRNAs (Mirg; ENSMUSG00000097391). The two secondary structure profiles show a strong a correlation (0.92).

Our results suggest that the secondary structure of RepA is conserved among primates, and diverges between human and mouse. However, analyzing the information contained in structured nucleotides (i.e., nucleotides with CROSS score < 0 are set to 0) we could identify *XIST* as the best match of human RepA within all mouse lncRNAs (structural distance 0.034; p-value < 10^−6^; **Figure 4C**). This result indicates that double-stranded regions are more conserved than single-stranded regions. In addition, we note that by sequence identity, *XIST* ranks as the 14^th^ hit of human RepA in all mouse lncRNAs, which indicates that methods based on sequence comparison show a significantly lower ability to identify structural homologues.

### HOTAIR

HOTAIR shows a complex secondary structure, divided into four domains (D1, 2, 3, 4; **Figure. 5A** and **5B**) [25]. Experimentally it has been determined that more than 50% of the nucleotides are involved in base pairing (CROSS achieves an AUC > 0.80 in predicting its SHAPE profile; **Supplementary Figure. 8A** and **8B**) [26]. The region D2 is highly structured and consists of 15 helices, 11 terminal loops, and 4 junctions (three 5-way junctions and one 3-way junction) [26].

**Figure 5.**
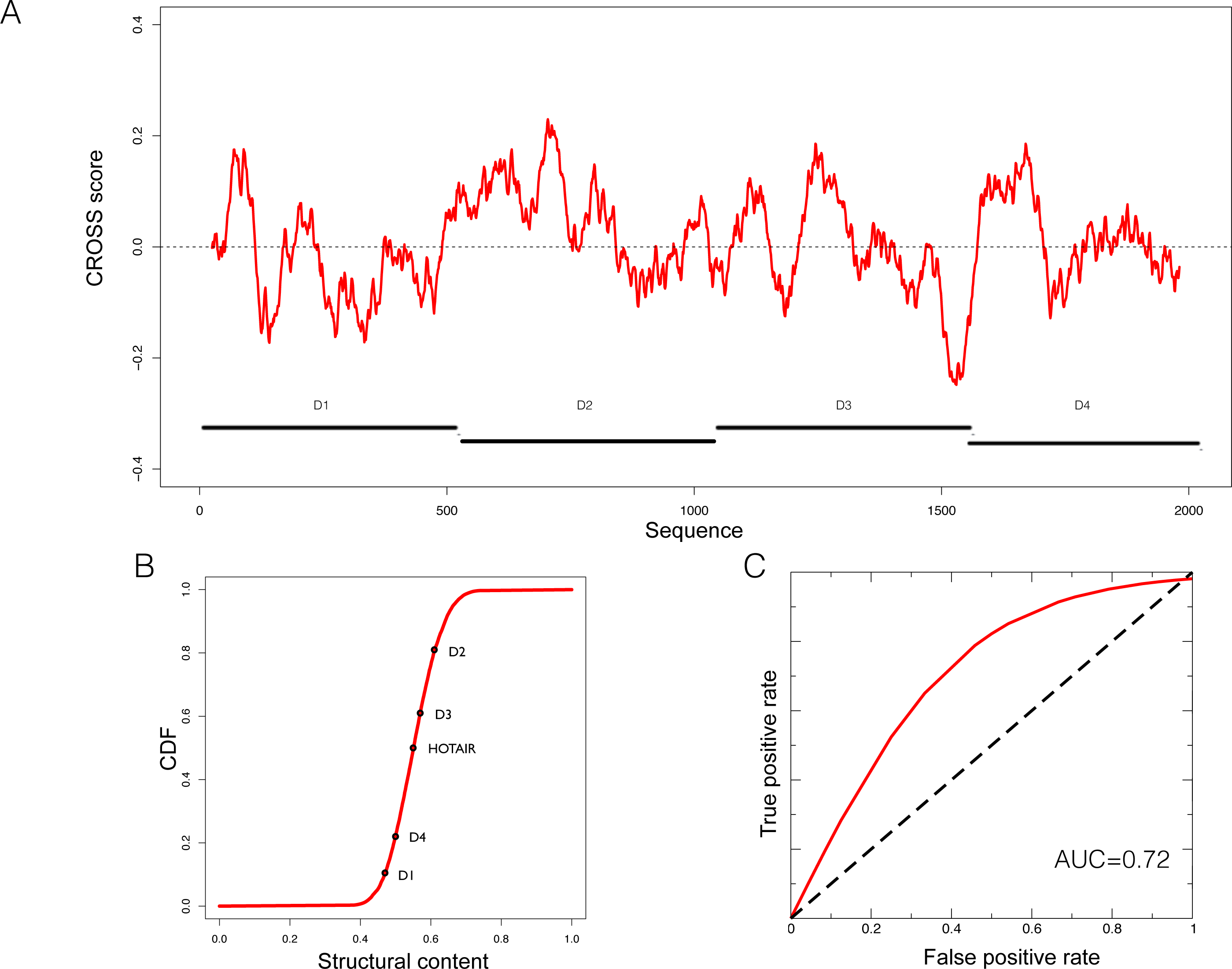
Predictions of HOTAIR structure. (**A**) Secondary structure profile of the complete murine *HOTAIR* obtained using CROSS. Positive regions are to be considered double-stranded, negative regions single-stranded. (**B**) Cumulative distribution function (**CDF**) of the structural content of all the human lincRNAs predicted by CROSS. The structural contents of the D domains of *HOTAIR* are reported on the curve. (**D**) ROC curve of *CROSSalign* to identify reverse engineered sequences with the same structure as D2 and a different sequence.

The D2 domain of *HOTAIR* is predicted by *CROSS* to be the most structured (**Figure 5B**). We used the same reverse engineering process as for RepA to generate 50 different sequences (average identity of 40%; **Supplementary Table 1**) with the same secondary structure as D2 (**Materials and Methods**; ***Reverse engineering: from structure to sequence***). Also in this case, *CROSSalign* reports good performances (AUC of 0.72; **Figure 5C**) that are superior to those obtained using the covariance-based approach *CMsearch* (AUC of 0.70; **Supplementary Figure 9**; **Materials and Methods**; ***Comparisons with CMsearch***) [20].

We then selected the D2 domain of HOTAIR to measure its conservation in 10 species [21] using *CROSSalign* (**Supplementary Figure 9A**). As for *XIST*, the structural distance analysis indicates that primates cluster close to human, and other species are more distant (**Supplementary Figure 9B**). Orangutan D2 was then searched for within all human lncRNAs, and HOTAIR was identified as the best match (structural distance 0.032; p-value < 10^−6^) with overlapping coordinates (nucleotides: 666-1191; 78% overlap with the query region; **Figure 6A**). Searching for mouse D2 within all human lncRNAs, HOTAIR was found as the best (0.092; p-value < 10-4) and matching position (nucleotides: 284-788; 57% overlap; **Figure 6B**). These results suggest that D2 secondary structure is not only conserved in primates but also in mouse.

**Figure 6.**
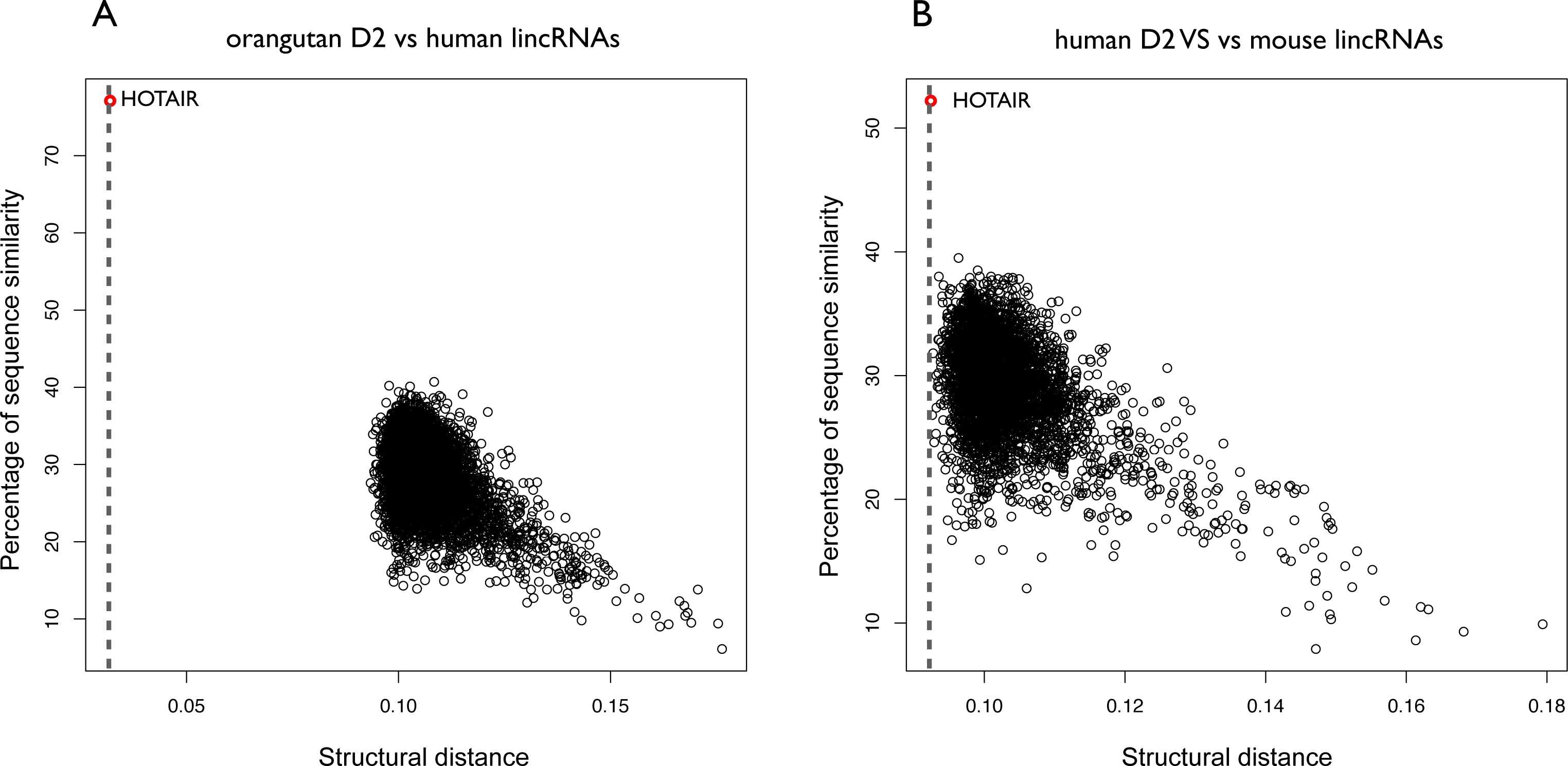
Structural conservation of HOTAIR D2 in different species. (**A**) Structural similarities of orangutan D2 against all human lincRNAs. The structural distance was obtained using OBE-DTW and plotted against with the sequence similarity. The D2 of orangutan is identified as the best match (colored in red). (**B**) Structural similarities of human D2 against all mouse lincRNAs. Human D2 is identified as the best match (colored in red).

To further investigate the secondary structure of HOTAIR, we studied the structural conservation of the D4 domain (**Supplementary Figure 10**). As opposed to D2, D4 is predicted by CROSS to be poorly structured (**Figure 5B**). Searching for orangutan D4 within all human lncRNAs yields HOTAIR as the best match (structural distance 0.023; p-value < 10^−6^) and the reported sequence position shows a sizeable overlap with the D4 domain in human HOTAIR (predicted coordinates: 1650-2291; overlap of 79%; **Figure 7A**). By contrast, when searching for mouse D4 within all human lncRNAs, HOTAIR shows poor ranking (1849^th^; structural distance 0.104; p-value = 0.061), which indicates little structural homology between human and mouse (**Figure 7B; Supplementary Figure 10**).

**Figure 7.**
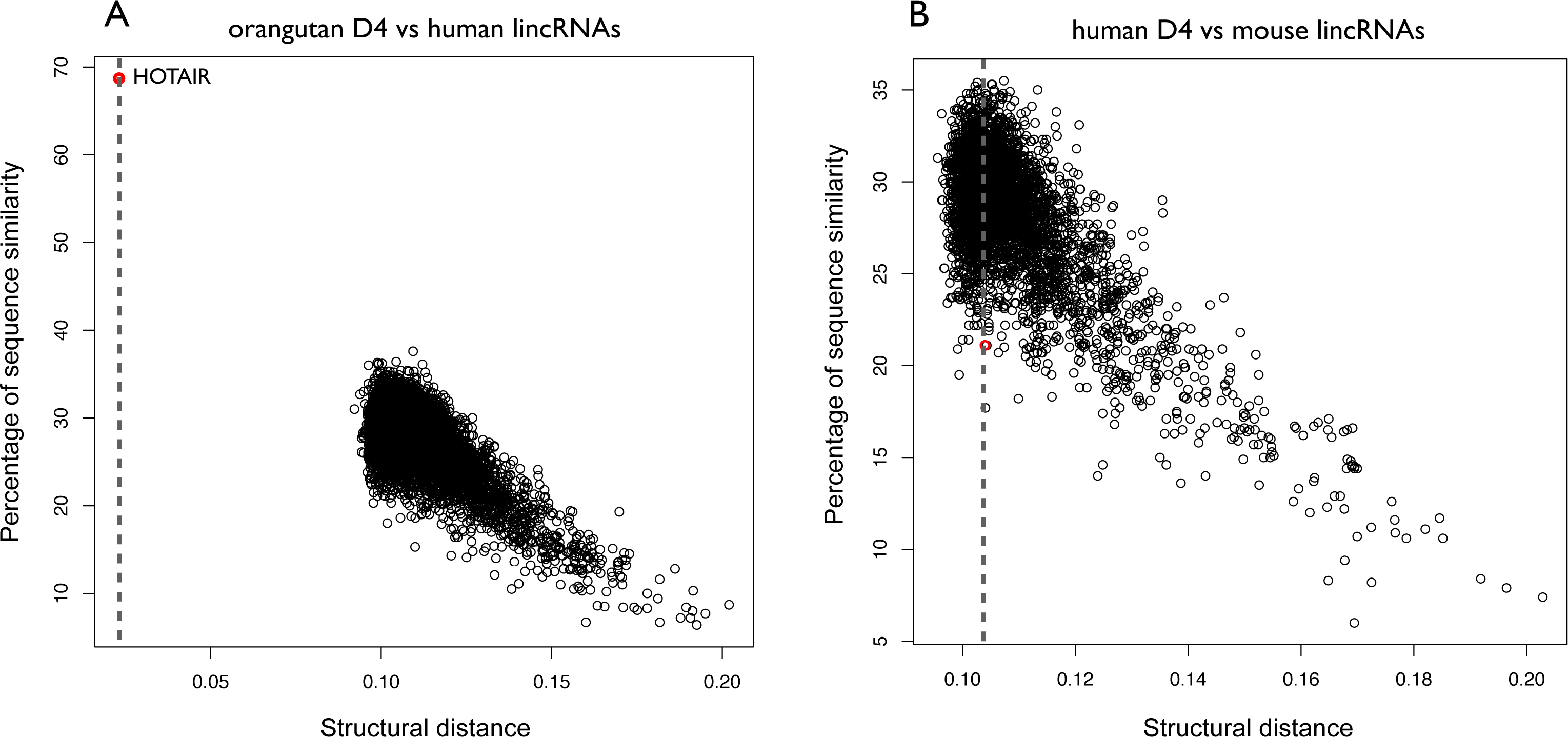
Structural conservation of HOTAIR D4 in different species. (**A**) Structural similarities of Orangutan D4 against all human lincRNAs. The structural distance was obtained using OBE-DTW and plotted against sequence similarity. Orangutan D4 is identified as the best match (colored in red). (**B**) Structural similarities of human against all mouse lincRNAs. Human D4 is not identified as the best match (1849th best hit; colored in red).

### HIV

HIV is one of the most studied ssRNA viruses with a complex secondary structure [27] that is accurately predicted by CROSS [10] (see also http://service.tartaglialab.com/static_files/algorithms/cross/documentation.html#4). As other organisms, HIV and ssRNA also contain non-coding regions [28].

We divided HIV into 10 non-overlapping regions of ~1000 nucleotides and searched each of them against a database of ssRNA viruses having as host human (292 cases, downloaded from NCBI; **Supplementary Table 3**) to identify structurally similar domains. We found that coronavirus HKU and Simian-Human immunodeficiency SIV have the most significant matches with HIV (structural distance 0.078, p-value < 10^−6^ for SIV; structural distance 0.093, p-value < 10^−4^ for HKU). This finding is particularly relevant since SIV and HIV share many similarities in terms of pathogenicity and evolution [29]. Indeed, previous studies already reported a similarity in terms of secondary structure between HIV and SIV that is not explained by sequence similarity [30].

In addition, we found that the HIV 5’ region is structurally similar to a strain of Ebola virus (Tai Forest; **Supplementary Table 3**). In agreement with this observation, previous studies indicate that HIV and Ebola have the same mechanisms of egress, taking contact with the cellular protein Tsg101 [31]. Moreover, HIV 5’ is the most conserved region in all ssRNA viruses (**Figure 8A**). This result indicates that the secondary structure of this region is not only necessary for HIV encapsidation [32], but is also essential for the activity of other viruses.

**Figure 8.**
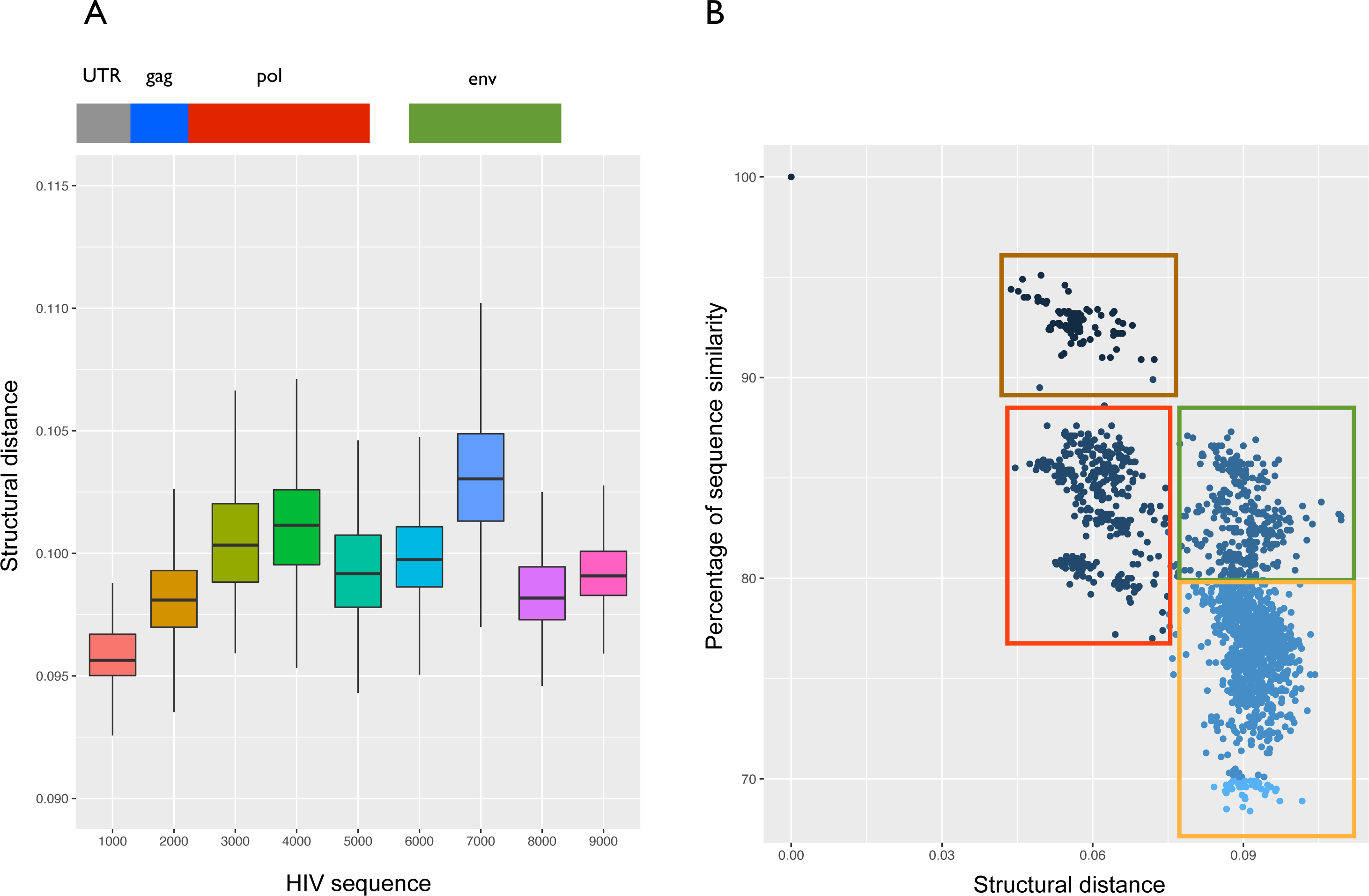
Structural analysis of the HIV transcriptome. (**A**) Structural conservation of HIV genome (divided into 10 not overlapping regions) compared with the complete genome of 292 ssRNA viruses. The region spanning the first 1000nt (including 5’ UTR) is the most conserved among all the viruses. (**B**) Structural distances of the complete HIV genome against the complete genomes of 4884 HIV strains. Using analysis of primary and secondary structures, we identified four main clusters (red, green, brown and yellow). Red and green boxes indicate strains whose structural difference cannot be identified through sequence analysis, while brown and red boxes as well as green and yellow boxes identify strains with similar structures and different sequences.

We also compared structural distances and sequence similarities of all HIV strains (4804; see **Materials and Methods**). We found two clusters (brown and red; **Figure 8B**) that are similar in terms of structure (~0.06 structural distance; p-value < 10^−6^) and sequence (80%-95% sequence similarity). Other clusters (red and green; **Figure 8B**) showed significant distance in structure (from 0.06 to 0.09 of structural distance; p-value < 10^−6^) that is not identifiable by sequence similarity (~85% sequence similarity). This result suggests that HIV could have evolved maintaining a similar sequence but different structures, as previously reported in literature [30].

## Discussion

We developed the *CROSSalign* method based on the combination of the CROSS algorithm to predict the RNA secondary structure at single-nucleotide resolution [10] and the Dynamic Time Warping (DTW) algorithm to align profiles of different lengths [11]. DTW has been previously applied in different fields, especially pattern recognition and data mining [34,35], but has never been used to investigate structural alignments of RNA molecules. Since CROSS has no sequence length restrictions and shows strong performances on both coding and non-coding RNAs [10] the combination with DTW allows very accurate comparisons of structural profiles. Thermodynamic approaches, such as RNAstructure [14] or RNAfold [12], cannot be directly used for such a task since they are restricted on the sequence length [36].

We applied *CROSSalign* to investigate the structural conservation of lncRNAs in different species and the complete genomes of ssRNA viruses. We found that the algorithm is able to find structural homologues between thousands of matches and correctly identifies the regions of similarity between profiles of different lengths. The results of our analysis reveal a structural conservation between known lncRNA domains including *XIST* RepA (best hit out of 8176 cases; 95% overlap with the query region) and *HOTAIR* D2 (best hit out of 8176 cases; 78% overlap with the query region), but also identify structural similarities between regulatory regions of HIV and other ssRNA viruses, opening new questions regarding similar mechanisms mediated by the secondary structure. Importantly, RepA and D2 profiles were accurately recognized in a pool of RNAs designed to have the same structure but different sequences. On these datasets multiple sequence alignments performed with *CMsearch* showed lower performances, indicating that *CROSSalign* predictions can be used to complement primary structure comparisons.

Our webserver is available at http://service.tartaglialab.com//new_submission/CROSSalign (documentation and tutorials are at the webpages http://service.tartaglialab.com/static_files/algorithms/CROSSalign/documentation.html and http://service.tartaglialab.com/static_files/algorithms/CROSSalign/tutorial.html) and allows to predict structural similarities between two (Standard, OBE, Fragmented modes) or more (Dataset, Custom dataset) RNA molecules. *CROSSalign* can be interrogated to search for structural similarities between thousands of lncRNA molecules and identifies regions with similar structures using a specific DTW algorithm (open begins and ends OBE).

As shown in the examples presented, *CROSSalign* is a versatile algorithm able to simplify the complex search for structural similarity among RNA molecules and shows great potential for the study of lncRNAs.

## Materials and Methods

### Prediction of the RNA secondary structure: CROSS

Secondary structure profiles were generated using CROSS [10]. CROSS models have been previously trained on data from high-throughput experiments (PARS: yeast and human transcriptomes [3,37] and icSHAPE: mouse transcriptome [38]) as well as on low-throughput SHAPE [27] and high-quality NMR/X-ray data [39]. Since each approach has practical limitations and a different range of applicability, we combined the five models into a single algorithm, Global Score, which provides a consensus prediction.

The consensus model *Global Score* was trained and tested on independent sets of NMR/X-ray structures (11’670 training fragments, 5’475 testing fragments [12,40], see also https://github.com/stanti/shapebenchmark/tree/master/benchmarkdata). In the testing phase, single and double-stranded nucleotides were recognized with an AUC of 0.72 and a PPV of 0.74. Comparison with experimental SHAPE data shows similar performances (AUC of 0.76 and PPV of 0.76; see http://service.tartaglialab.com/staticfiles/algorithms/cross/documentation.html#5) and all the details are reported in our previous publication [10]. In addition, as done with experimental SHAPE data, Global Score has been also used as a constraint in RNAstructure [13,14]. On the test set [12], Global Score was shown to increase the PPV of RNAstructure from 0.68 to 0.72, with remarkable improvements in 13 cases (from 0.44 to 0.72) and decreases the PPV in only three cases for which real SHAPE data does not improve performances. Moreover, using the partition function computed with RNAstructure, we previously calculated the AUC for each structure with and without CROSS constraints and observed an improvement from 0.81 to 0.86 when CROSS is integrated in the algorithm. We observed a similar trend using RNAfold [12] (the PPV increases from 0.67 to 0.70 using Global Score and the AUC remains at 0.85).

In the present study all the profiles were computed using the *Global Score* module: nucleotides with a score higher than 0 are predicted to be double-stranded and structured, while nucleotides with a score lower than 0 are single-stranded. Since the algorithm has no sequence length restriction and shows strong performances on both coding and non-coding RNAs [10] it was combined with DTW for pairwise comparison of structural profiles. Thermodynamic approaches, such as RNAstructure [14] *or RNAfold* [12], could not be directly used for such task since they are restricted on the sequence length.

### Comparison of structural profiles: DTW

To compare two CROSS profiles, we used the Dynamic Time Warping (*DTW*) algorithm available in the R package dtw [11]. The open begin and end (*OBE-DTW*) algorithm was employed to compare profiles of different lengths. Indeed, the standard *DTW* method imposes the same begins and ends to the two profiles that are compared, while *OBE-DTW* searches for the profile of shorter length within the other one. Accordingly, we used standard *DTW* to compare profiles of similar lengths (i.e., one sequence is less than 3 times longer than the other), while *OBE-DTW* was employed to search for modules within larger profiles (e.g., RepA within the complete *XIST* sequence; *XIST* is ~45 times longer than RepA).

The structural distance is computed with an asymmetric pattern and using the Manhattan distance, which is optimal for comparing profiles of different lengths. To avoid biases regarding the length of the profiles, the final structural distance is normalized for the lengths of both profiles using the internal function *normalizedDistance*. We also tested different normalizations of DTW outputs (including normalization by length of the shorter or longer profile) and we found that the normalization based on the lengths of both profiles is optimal. The function *index* was used to visualize the optimal path and to extract the matching coordinates between the two profiles.

### Statistical analysis

To compute the significance of a specific DTW score, we analyzed the statistical distributions generated using human lncRNAs of different lengths (200, 500, 1000, 5000 nucleotides). 100 molecules for each class were employed to compute the structural distance between the classes. The distributions are set as a reference to compute the p-values in new analyses (**Supplementary Table 4**)

### Datasets

- lncRNA sequences were downloaded from ENSEMBL 82 using Biomart and specifying lincRNAs, for a total of 4427 sequences for mouse and 8176 for human.
- The complete viral genomes were downloaded from NCBI selecting ssRNA viruses having as host human or primates (for SIV), for a total of 292 complete genomes.
- The complete rRNA sequences were downloaded from NCBI.
- RepA, D2 and D4 were selected from the data publicly available from the work of Rivas et al., 2016 [21]. To keep consistency between the results we tried to select the same species between the two sets of multialignments. When this was not possible we selected similar species (rat and mouse, orangutan and chimpanzee, lemur and sloth).
- The HIV strains were downloaded from HIV databases (https://www.hiv.lanl.gov/), selecting only complete genomes for a total of 4804 sequences processable by CROSS.

### Sequence alignment

To compute the sequence alignments we used the browser version of EMBOSS-needleall, publicly available at http://www.bioinformatics.nl/cgi-bin/emboss/needleall. The tool was used with standard settings to speed up the calculation. The sequence identity was retrieved from the corresponding field from EMBOSS multiple output.

### Reverse engineering: from structure to sequence

To study how sequence similarity is related to structural similarity we created different sequences with the same secondary structure as RepA (*XIST*) and D2 (*HOTAIR*). To generate the reference structure we used *RNAfold* [41]. We then generated different sequences encoding for the same previously generated structure. For this task we used the command line version *RNAinverse* from the Vienna suite [12]. The tool was launched using standard parameters to generate 50 sequences for each structure.

### Comparisons with CMfinder

We compared the structural distances provided by *CROSSalign* with the multiple sequence alignment scores of CMfinder [15]. For this analysis we selected 5 of the most complex datasets (i.e. highest number and longest sequences) from the test set of *CMfinder* (cobalamin, intron gp II, sbox, lysine and histone 3). To compare the pairwise distances (*CROSSalign*) with the multiple alignment scores (CMfinder) we computed all the distances within the datasets and selected the lowest distance for each transcript. From low-(median) to high-confidence (top and bottom 5%) CMfinder scores, we observed an increase in the performances of *CROSSalign*, which indicates a good predictive power on the multiple alignment score.

### Comparisons with CMsearch

*CMsearch* is a method used to search a covariance model (CM) against a sequence database and provides a ranked list of the sequences with the most significant matches relative to the CM. Using the CMbuild package we built the CMs using as input the multiple alignments of the two reference sets (RepA and D2). The E-values of the CMs were obtained upon calibration with *CMcalibrate*. The calibrated CMs were used to search for homologues in the positive and negative sets using the *CMsearch* approach. By running *RNAalifold*[42] to build a consensus secondary structure on aligned sequences[43], we did not obtain improvements for both RepA and D2 alignments.

### Algorithm Description

#### Input

The user should paste one or two RNA sequences in FASTA format into the dedicated form, providing an email address (optional) to receive a notification when the job is completed. The algorithm can be launched in 4 different modes, each of them being a specific variation of the DTW algorithm (**Supplementary Figure 1**).

- The standard-DTW is recommended for comparing structures of RNAs with similar lengths (i.e., one sequence is less than 3 times longer than the other).
- *OBE-DTW* (open begins and ends) is a specific mode to search for a shorter profile within a longer one. This is the recommended mode when comparing profiles of very different sizes (i.e., one sequence is more than 3 times longer than the other). Please keep in mind that the sequence in the form of RNA input 1 will be searched for within RNA input 2, so the sequence in RNA input 1 should be shorter than the other.
- The fragmented OBE-DTW is a specific mode for searching for unknown secondary structure domains of one profile within the other. The secondary structure of RNA input 1 will be fragmented with a non-overlapping window of 200 nucleotides [optimal size to search secondary structure domains in large RNAs [44,45]] Each fragment of RNA input 1 will then be searched for within the other sequence. This approach is the recommended mode when the user is not interested in the global similarity between two secondary structure profiles, but wants to search for an unknown domain conserved in both sequences. A minimum length of 600 nucleotides is recommended for fragmentation.
- The *dataset* mode allows the user to search for a single sequence within all the lincRNAs of a specific organism. The shorter profile of each pair will be searched for within the larger one following the *OBE-DTW* procedure. The organisms available are Human, Mouse, Rat, Macaque and Zebrafish. The lincRNAs where downloaded using Biomart (Ensemble 82). New organisms and updated versions will be regularly added.

#### Output

We report the *CROSSalign* score that measures the structural distance between two structures. The closer the score is to 0, the higher the similarity in terms of secondary structure. According to our statistical analysis, RNA molecules with a structural distance of 0.10 or higher are to be considered different in terms of secondary structure (see online ***Documentation***).

The main image shows the overall structural similarity of the two profiles employed to calculate *CROSSalign* score (**Supplementary Figure 2A**). On the two axes the user will see the structural profiles obtained with CROSS for the two RNA sequences in input (score >0 means a double-stranded nucleotide; <0 single-stranded;). For a better visualization, the profiles are smoothed using a function previously defined [10]

The similarity is represented by the red path in the figure, obtained with the index function of the dtw package. The closer the path is to the diagonal, the more similar are the profiles. Vertical or horizontal paths are to be considered gaps, while diagonal paths highlight similar regions of the two profiles.

Since *OBE-DTW* allows the identification of the optimal starting/ending points of a match, the optimal match is reported in terms of coordinates relative to the larger profile (RNA input 2). The main plot shows the CROSS profiles of the optimal matching region selected by the *OBE-DTW* algorithm (**Supplementary Figure 2B**). In order to keep the gaps introduced by the *OBE-DTW* algorithm, the two profiles are not smoothed.

The *fragmented OBE-DTW* is a particular form of *DTW* optimized to search for all the possible structural domains of a particular sequence within another one. The main output is a scrolling table reporting the structural score, the beginning of the match, the end of the match and the p-value (**Supplementary Figure. 2C**). All the values are computed with the procedure used for *OBE-DTW*. The table can also be downloaded as a .txt file. The same output is used for the dataset mode, but in this case the table can only be downloaded.

## ACKNOWLEDGMENTS

We thank Fernando Cid and the members of Tartaglia’s group for useful comments.

The research leading to these results has received funding from the European Union Seventh Framework Programme (FP7/2007-2013), through the European Research Council, under grant agreement RIBOMYLOME_309545 (Gian Gaetano Tartaglia), and from the Spanish Ministry of Economy and Competitiveness (BFU2014-55054-P and BFU2017-86970-P). We also acknowledge support from AGAUR (2014 SGR 00685), the Spanish Ministry of Economy and Competitiveness, ‘Centro de Excelencia Severo Ochoa 2013–2017’ (SEV-2012-0208). We also thank the CRG fellowship to SM.

## AUTHORS CONTRIBUTIONS

GGT designed the work, RDP implemented the approach with SM. AA and RDP developed the server. GGT, RDP and SM wrote the paper. All authors reviewed the manuscript.

## SUPPLEMENTARY MATERIAL

See accompanying document.

## AVAILABILITY OF DATA

The code is publicly available under an open source license compliant with Open Source Initiative at https://github.com/armaos/algorithm-CROSSalign. The source code is deposited in a DOI-assigning repository https://doi.org/10.5281/zenodo.1168294.

